# The amplitude and latency of the earliest signal in V1 encode bottom-up saliency by feature conjunction

**DOI:** 10.64898/2026.01.17.699936

**Authors:** Chen Wu, Xiaoning Li, Huan Li, Xuan Wang, Ziang Yin, Zeyu Wang, Peng Zhang, Zhikuan Yang, Jinyou Zou

**Author notes:** **Correspondence to:** Z.Yang, J.Zou.

## Abstract

The neural origin of bottom-up saliency for exogenous attention remains highly controversial. In this study, we investigated whether the earliest activity in the primary visual cortex (V1) encodes saliency signals defined by the eye-of-origin and feature-conjunction information. Electroencephalography (EEG) recordings from the human occipital cortex revealed early responses to eye-of-origin (E) and/or orientation (O) singletons, with larger response amplitudes to the double-feature (EO) singletons. The short onset latency (58–70 ms) and polarity reversal of the responses indicate a V1 origin. Importantly, the latency and amplitude of these responses predicted behavioral detection performance. Together, these findings suggest that the timing and amplitude of the earliest signals in V1 represent the saliency of combined feature contrasts for bottom-up attention. These signals unlikely originate from projections of other proposed source areas of saliency, due to the scarcity of necessary monocular neurons to process eye-of-origin information.

**Highlights:**

- Eye-of-origin information, invisible to the SC, elicits an early saliency signal in V1 within 50–100 ms.
- Combined feature contrast enhances V1 saliency responses in a nonlinear fashion.
- The latency and amplitude of V1 saliency responses predict behavioral detection performance.

## Introduction

Visual attention selects limited inputs from an overwhelming amount of sensory data via top-down (endogenous) and bottom-up (exogenous) mechanisms (Corbetta and Shulman, 2002). Exogenous attention rapidly identifies salient features, such as unique colors or orientations (Wolfe, 2021), acting as a first-line gatekeeper that enables automatic extraction of evolutionarily critical information (Zhaoping, 2023). However, its neural origin remains debatable. While early research implicated high-level cortical areas such as the intraparietal sulcus (IPS) (Gottlieb *et al*., 1998), recent converging evidence supported the V1 Saliency Hypothesis (V1SH) which proposed that the primary visual cortex (V1) generates saliency maps indicating attractiveness at different visual locations to guide exogenous attention (Li, 1999). Conversely, primate electrophysiology suggests that the superior colliculus (SC) encodes saliency before V1 does (Veale *et al*., 2017; White *et al*., 2017). The roles of the SC and V1 in visual saliency have been a bone of contention among researchers. Thus, these conflicting perspectives need to be further explored. For clarity, in this paper we use the term “saliency” to describe specifically the strength of bottom-up attraction of unique features among distracting items (Itti and Koch, 2001; Nothdurft, 2000; Treisman and Gelade, 1980).

The V1SH is supported by numerous behavioral observations, particularly those regarding the prediction about a specific feature, the eye of origin—that is, whether an object is input from the left or right eye; it is more broadly referred to as the difference in input strength between the two eyes. The V1SH predicts that eye-of-origin contrasts, such as an object (an eye-of-origin singleton) that presents in the left eye while all others present in the right eye, captures attention even if it is typically elusive to conscious perception (Ono and Barbeito, 1985). Empirical data confirmed this prediction by showing attention attraction by eye-of-origin singletons in a visual search task (Zhaoping, 2008, 2012). Since the monocular neurons necessary for the computation of eye-of-origin contrasts are abundant in V1 (Dougherty *et al*., 2019) but scarce in SC (Finlay *et al*., 1976b; Schiller *et al*., 1974) and higher cortices (Zhaoping, 2008), this effect is believed to be a hallmark of V1 mechanisms. Direct neural evidence of attentional capture by eye-of-origin singletons could be critical for comparing the V1SH with competing theories.

Zhaoping (2016) proposed that saliency computation has migrated from SC in lower vertebrates to V1 in primates with evolution. According to the V1SH, the response strength of feature-tuned V1 neurons represents visual saliency. Neurons mutually inhibit neighbors tuned to similar features, thus making responses to unique features “pop out” in the saliency map, which is realized by the SC for subsequent actions. Studies that utilized electroencephalography (EEG) and functional magnetic resonance imaging (fMRI) confirmed that V1 encodes saliency driven by orientation contrasts (e.g., a horizontal bar surrounded by vertical bars) within 50–100 ms (Liu *et al*., 2025; Zhang *et al*., 2012b). However, contradictory results were found in electrophysiological research on monkeys: While Yan et al. (2018) supported the observations in humans, White et al. (2017) reported that the saliency representation in V1 emerges after 100 ms and after the representation in SCs, providing strong evidence against V1SH.

Here, we further investigated the relationship between V1 and the SC by recording the cortical activity associated with attention driven by eye-of-origin contrasts. As the eye of origin cannot be realized by binocular neurons in the SC, the results reflect intracortical mechanisms with minimal tectostriatal influence. If saliency signals from feature contrasts are first decoded in the SC and then sent back to the cortex, early cortical saliency responses to eye-of-origin singletons may be absent or substantially different from responses to other feature (e.g., orientation) singletons. Conversely, if saliency signals are decoded during V1 feedforward processing, saliency responses should be observed in V1 within 100 ms. These early responses should also be related to the increased salience resulting from the combination of multiple features (Koene and Zhaoping, 2007; Krummenacher *et al*., 2001; Treisman and Gelade, 1980; Zhaoping, 2008; Zhaoping and Zhe, 2015). To test these predictions, we recorded and compared EEG responses specific to eye-of-origin and/or orientation singletons in human observers. Our analysis focused on early V1 responses within 100 ms and their correlation with behavioral performance. These results will provide valuable insights into the role of V1 mechanisms in saliency processing.

## Results

### Uniqueness in the eye of origin captures visual attention

We first confirmed the attentional effect of eye-of-origin contrasts in behavioral tasks. Participants were asked to maintain central fixation and identify a unique line at an eccentricity of 9° (Figure 1A). In different conditions, the target is a feature singleton unique from all other lines in either eye of origin (E), orientation (O), eye of origin and orientation conjunctively/redundantly (EO). Stimuli were only presented for 50 ms to minimize confounding from eye movements. Subjects made a forced choice to indicate, as accurately as possible, whether the target appeared on the left or right half of the screen via the press of a button (Figure 1B).

**Figure 1.**
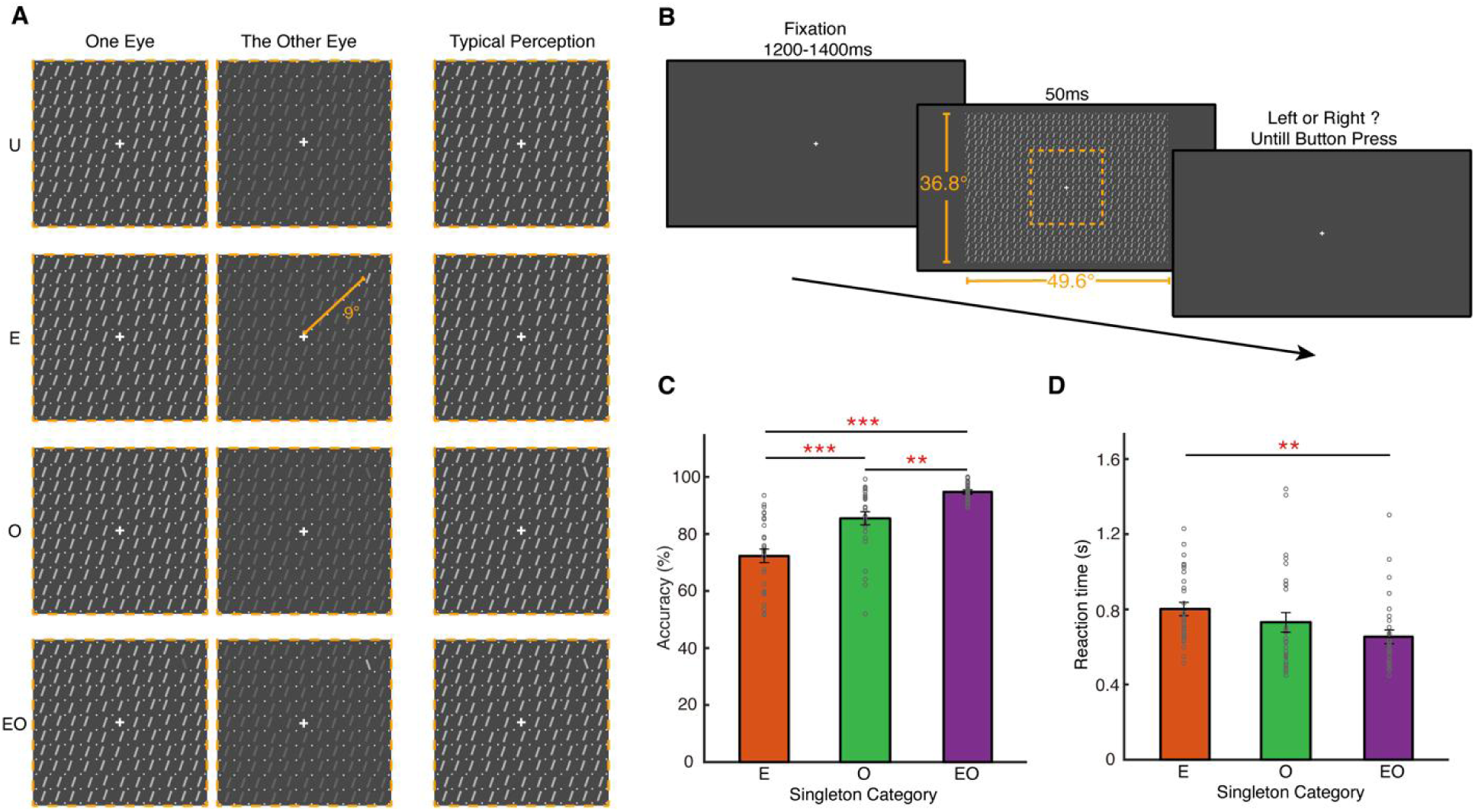
Stimulus and Behavioral Performance. **(A)** Visual stimulus. In different conditions, either an eye-of-origin (E) singleton, an orientation (O) singleton, an eye-of-origin and orientation double-feature (EO) singleton, or no singleton (U) was presented in periphery (at an eccentricity of 9°) among a dense array of bars. For illustration purposes, only bars in the center of the screen are depicted in this figure. **(B)** Behavioral experiment procedure. Each trial began with a 1200–1400 ms fixation, followed by the bar stimulus for 50 ms. Subjects pressed one of two buttons to indicate whether the singleton appeared on the left or right of the screen. The dashed box (not shown in the experiment) indicates the stimulus region depicted in (A). **(C)** Accuracy and **(D)** reaction times for the behavioral tasks. Gray circles indicate the data of individual subjects; bars represent group means ± SEM. ** and *** indicate Holm-corrected p < 0.01 and p < 0.001.

One-way repeated-measures ANOVA revealed a significant effect of singleton category on task accuracy (F(2, 56) = 30.055, p < 0.001, 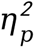 = 0.518). Post-hoc tests revealed that the performance in the EO condition was significantly better than that in the O condition (t(28) = 2.890, p = 0.005, Cohen’s d = 0.664), thereby indicating a saliency effect through additional uniqueness in eye of origin. Further, the performance in the E condition was much lower than that in the EO (t(28) = 7.676, p < 0.001, d = 1.765) and O conditions (t(28) = 4.785, p < 0.001, d = 1.100), thus revealing the difficulty in recognizing E singletons (Figure 1C). These results were further supported by a significant effect (F(2, 56) = 6.449, p = 0.003, 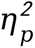 = 0.187) of singleton category on reaction times (RTs). RTs decreased from the E condition to the O condition to the EO condition, though only the difference between the E and EO conditions reached a significant level (post hoc t(28) = 3.590, p = 0.002, d = 0.664, Figure 1D). Additionally, two-way repeated-measures ANOVA involving both factors of singleton categories and target locations (upper vs. lower visual fields: Figure S1; or nasal vs. temporal visual fields: Figure S2) revealed no significant main effect or interaction of target locations on response accuracy or RTs.

### Uniqueness in eye of origin and/or orientation induces early effects in occipital responses

In the EEG experiment, we first located the electrodes of interest, best representing early V1 responses, in an independent localizer session with a single star object (without surrounding objects) as visual stimulus. Six occipital electrodes with, on average, the strongest C1 troughs for UVF stimulation and the strongest C1 peaks for LVF stimulation were selected, including E33 (PO3), E39 (O2), E34 (PZ), E36 (POZ), E37 (OZ), E38 (PO4) (Figures 2A and 2B). In the main EEG session, we adopted similar visual stimuli of feature singletons as in the behavioral test, but with a different task. Participants maintained central fixation and passively viewed the bar stimulus, which was presented for 50 ms in each trial. To help maintain eye gaze, they responded to an occasional fixation color change that did not overlap with the timing of the bar stimulus (Figure 2C). This setup minimized the potential impact of top-down modulation and eye movement artefacts on the recorded neural activity related to exogenous attention. The results were then averaged over the electrodes of interest to obtain the event-related potentials (ERP).

**Figure 2.**
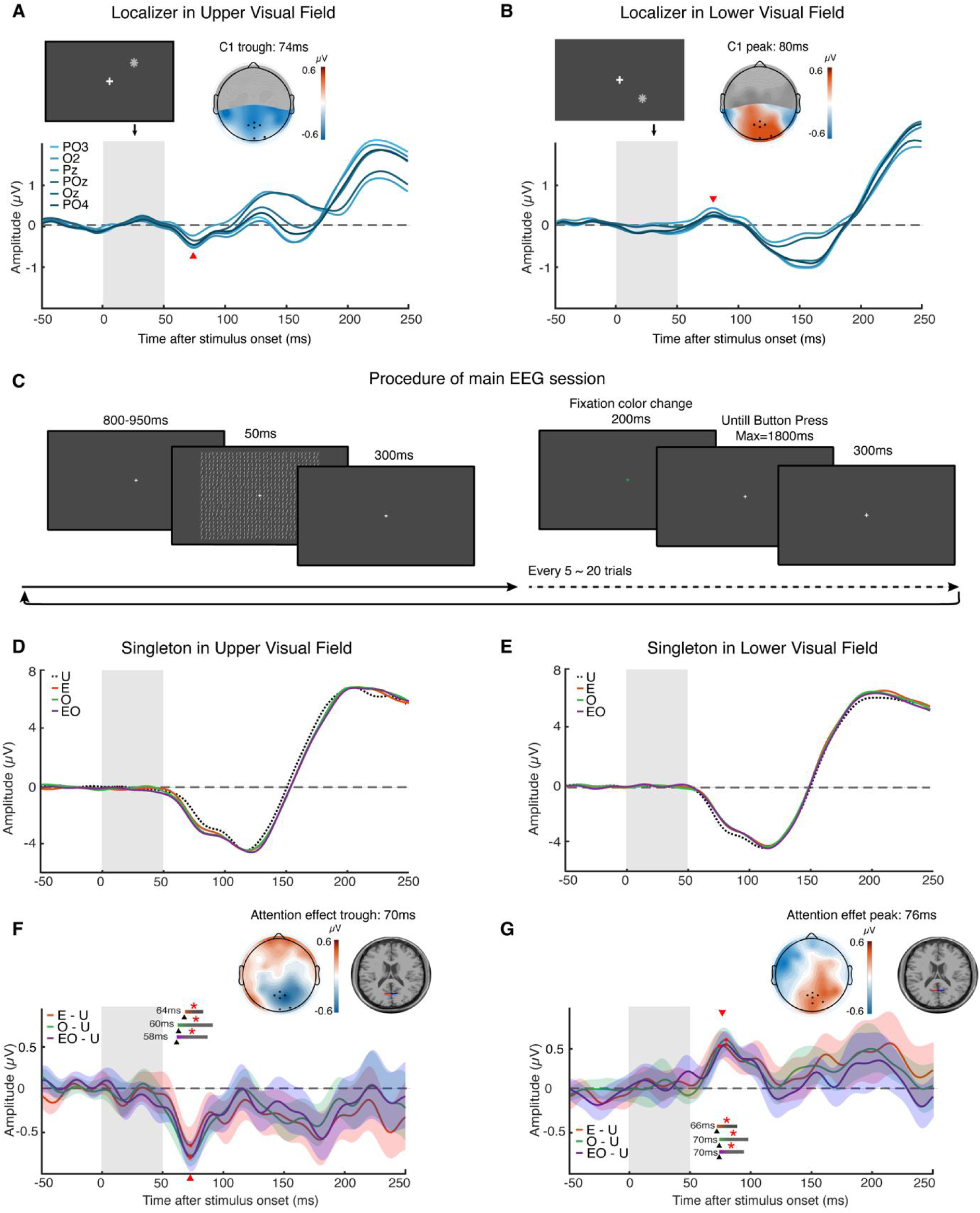
Procedures of and Results from the EEG experiments. **(A, B)** Electrode localizer session. The stimulus was a single star-shaped object present at one of the singleton locations in the main EEG sessions. The curves depict the average ERP across subjects, on each of the six electrodes with the strongest C1 amplitude. The vertical gray bars indicate the time interval of stimulus presentation. Topographic maps were obtained at the time of C1 peak/trough (red arrows). Black dots indicate the location of the selected electrodes among all the occipital electrodes (highlight regions). **(C)** Protocol of the main EEG session. Subjects passively viewed the bar stimulus and responded to occasional fixation color change. **(D, E)** ERPs from the main EEG session. Curves represent the average ERP across all subjects and the six electrodes of interest, for each condition separately. **(F, G)** Singleton-specific effects in the main EEG session. Curves display the response difference between the singleton-present conditions and the baseline U condition. Colored shadings denote standard error of the mean (SEM) across subjects. Horizontal bars represent the time intervals when the response was significantly different from the baseline (*for p < 0.05 in cluster-size based permutation test with additional Holm-correction). Black arrows and numbers indicate the starting time of the significant interval. Topographic maps and dipole source were shown based on the average responses of the three singleton-present conditions at the peak/trough time of the effect (red arrows).

In the ERP waveform, there appeared to be two negative components before and after 100 ms, respectively, although these were not clearly isolated. This general waveform was similar for singletons in the UVF and LVF (Figures 2D and 2E), and did not resemble the C1 polarity reversal response patterns. This was most likely due to the use of whole-screen stimuli (unlike the localizer sessions), which simultaneously activate the upper and lower parts of V1 subregions asymmetrically located in the calcarine sulcus (see Discussion). To further reveal the specific responses to the feature singletons, the ERP in a baseline U condition (singleton-absent condition, Figure 1A) was subtracted from that in the E, O, and EO conditions (singleton-present conditions). When the singleton appeared in the UVF, specific responses were found for E (64–78 ms, cluster p = 0.05), O (60–88 ms, cluster p = 0.049), and EO (58–82 ms, cluster p = 0.049) conditions, with an effect trough at 70 ms from stimulus onset in all three conditions (Figure 2F). When the singleton appeared in the LVF, specific responses were also found for E (66–88 ms, cluster p = 0.026), O (70–100 ms, cluster p = 0.034) and EO (70–94 ms, cluster p = 0.034) conditions, with an effect peak at 74 ms, 78 ms, and 78 ms, respectively (Figure 2G).

These responses had a rather short latency close to that of the first component in the general ERP waveform (Figures 2D and 2E) and the C1 component observed in the localizer session (Figures 2A and 2B), thereby implying a V1 source. Topographic maps confirmed that the neural origin of the observed effects was the occipital cortex. Further dipole source estimation localized symmetrical pairs of dipoles at V1 (Talairach coordinates: ±1, −69, and 4 for singletons in the UVF, and ±1, −52, and 2 for singletons in the LVF), accounting for over 90% of the variance in the scalp voltage at the trough (70 ms)/peak (76 ms) time of the effects.

### Polarity of the early EEG effects reverses according to the vertical position of singletons

In the results presented above, the singleton-specific effect in the ERPs was negative for singletons in the UVF and positive for singletons in the LVF, thereby exhibiting a prominent reversal of polarity depending on the vertical location of the singleton (Figures 2F and 2G). This is a unique characteristic of EEG signals from V1, and this is due to its anatomical organization in the calcarine sulcus, thereby strongly suggesting its V1 sources.

To further verify the robustness of the effect polarity, we identified the largest trough for UVF singletons and the largest peak for LVF singletons, within a 20–100 ms window for each participant. We calculated the peak-to-trough difference as the effect value of combining the UVF and LVF (Figure 3). We then compared this value with the result obtained using the same procedure on polarity-reversed EEG data (equivalent to finding the peak/trough for the singleton in the UVF/LVF using the original non-reversed EEG signals). Compared with the polarity-reversed data, the original (non-reversed) EEG signals showed significantly stronger peak-to-trough difference in all E (t(28) = 2.906, p = 0.007, d = 0.54), O (t(28) = 3.388, p = 0.004, d = 0.629), and EO (t(28) = 5.468, p < 0.001, d = 1.015) conditions. Furthermore, these values were tested in a permutation test and were found significant for all conditions (E: p = 0.006; O: p < 0.001; EO: p < 0.001). These results suggest that the observed EEG effect and its polarity reversal originate from true physiological activities in V1 rather than random noise of EEG signals.

**Figure 3.**
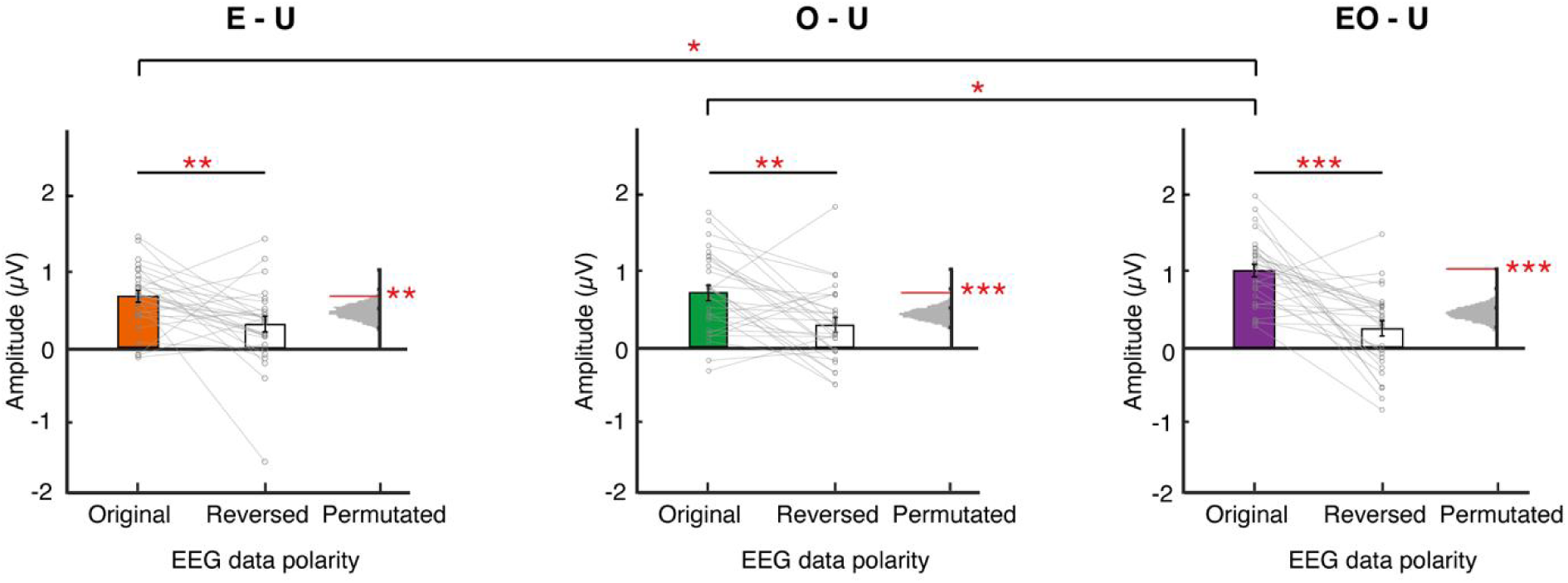
Singleton-Specific Effects in EEG Responses. Gray dots depict the amplitude of the singleton-specific effect (singleton-present condition vs. baseline condition) detected in each subject’s data. The effect was combined in the upper and lower visual fields by subtracting the trough in the upper-visual-field conditions from the peak in the lower-visual-field conditions (peak-to-trough difference). Bars indicate the mean value and SEM across subjects. The color-filled and empty bars depict results from original EEG data and polarity-reversed EEG data, respectively. Gray histograms indicate the null distributions calculated from the polarity-randomized EEG data in permutation tests. P-values are obtained as the position (red lines) of the original results in the null distribution. *, **, and *** represent Holm-corrected p < 0.05, p < 0.01, and p < 0.001.

We also tested whether the singleton category would affect this effect, especially when considering double-feature conditions versus single-feature conditions. One-way repeated-measures ANOVA revealed a significant effect of singleton category on the peak-to-trough difference (F(2,56) = 4.208, p = 0.02, 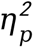 = 0.131). Post-hoc tests indicated that the EO condition elicited a significantly larger effect than both the O (t = –2.335, p = 0.046, d = 0.609) and E (t = –2.658, p = 0.031, d = 0.693) conditions (Figure 3). However, no significant effect of singleton category was found for the peak latency of the effect (p = 0.625). These findings suggest that combining multiple feature contrasts could strengthen the early attention-related response in V1.

To examine if the observed EEG effect was confounded by potential saccades toward the singletons, we performed a control analysis of the vertical electrooculograms (VEOG). The results showed no significant VEOG responses specific to singletons either in the UVF or LVF, within 200 ms after stimulus onset (uncorrected p > 0.05 for all time points). Since upward and downward saccades can generate opposite-polarity artifacts in the reference mastoid electrodes (Plöchl *et al*., 2012), we also examined the responses in these electrodes and found no singleton-specific effects. Thus, these analyses excluded potential influences due to eye movements (Figure S3). Moreover, we examined whether the observed effect in the E and EO conditions could be explained by a luminance disparity between the stimuli presented to each eye, potentially resulting from calibration errors. To investigate this possibility, we compared the C1 amplitudes elicited by left- and right-eye stimuli in the localizer session. The results showed no significant difference at each test visual location (Figure S4A). Psychophysically, subjects may perceive images as slightly brighter in the dominant eye than in the non-dominant eye, even when the physical inputs are matched between the two eyes (Porac and Coren, 1984). Therefore, we also performed the above analysis with the data grouped by dominant eye and non-dominant eye. The results showed no significant difference either (Figure S4B). These analyses thus suggested that the observed EEG effect was not an artifact of luminance differences between the two eyes.

### The early EEG effects correlate with behavioral performance

Finally, we tested whether the neural responses were associated with performance on the singleton-detection task among different subjects (Figure 4). First, we found that the response accuracy was positively correlated with the amplitude of the observed EEG effect (i.e., the peak-to-trough difference) in the pooled E, O and EO conditions (r = 0.453, p < 0.001), as well as in each singleton condition. In contrast, no significant correlation was observed between the response accuracy and EEG effect latency (uncorrected p = 0.315). Second, behavioral RTs were significantly correlated with both the amplitude (r = –0.398, p < 0.001) and latency (r = 0.474, p < 0.001) of the EEG effect. The latency appeared to be more closely related to the RTs.

**Figure 4.**
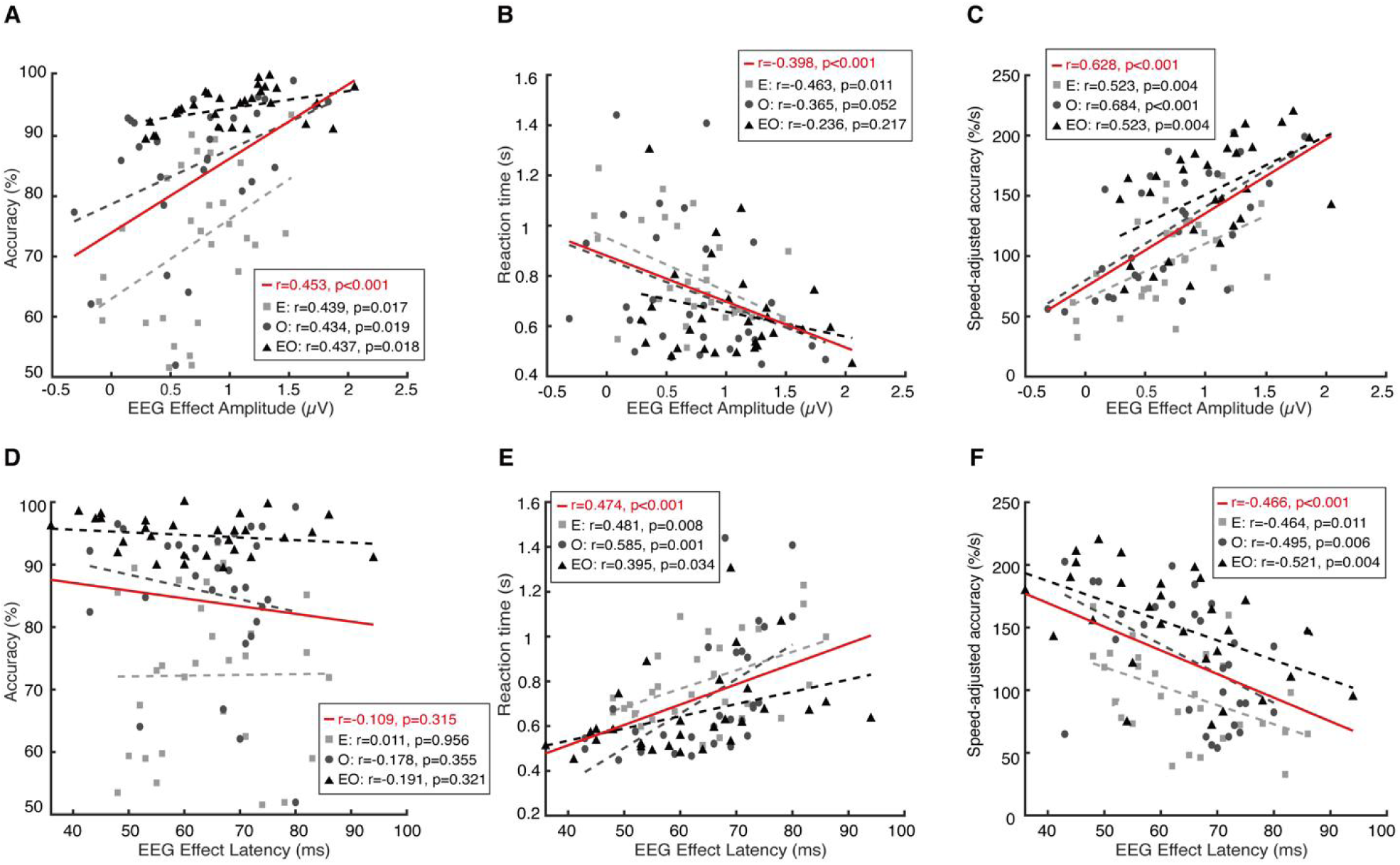
Correlation Between Neural Responses and Behaviors. Correlations between the early singleton-specific effect in EEG responses **(A-C**: amplitude; **D-F**: latency) and singleton-detection performance **(A, D**: accuracy; **B, E**: reaction time; **C, F**: speed-adjusted accuracy). Light gray, dark gray, and black denote data from the E, O, and EO conditions, respectively. Red lines indicate the overall correlation across all conditions. The p values for the overall correlations were corrected for multiple comparisons. Those for each singleton condition were not corrected.

Third, we calculated speed-adjusted accuracy, defined as response accuracy normalized by RT (similar to rate-correct score (Woltz and Was, 2006) proposed before), to better reflect behavioral efficiency that integrated both accuracy and RT measures. We found significant correlations between speed-adjusted accuracy and the amplitude (r = 0.628, p < 0.001) and latency (r = –0.466, p < 0.001) of the EEG effect. These results demonstrate a close relationship between behavioral performance, particularly detection efficiency, and very early occipital neural responses, therefore, suggesting that the V1 feedforward response represents exogenous attention. Furthermore, neural response amplitude and latency may primarily influence detection accuracy and speed, respectively, and the amplitude may also affect detection speed.

## Discussion

Our findings revealed early occipital responses to either eye-of-origin or orientation feature singletons. These responses occurred between 50 and 100 ms and exhibited a robust polarity reversal characteristic according to the vertical position of the singleton. Furthermore, these neural signals were closely linked to the singleton detection performance. Specifically, the conjunction of two visual features (EO singletons) enhanced the strength of early neural responses and facilitated the detection performance. Between-subject correlations suggest links between neural latency and behavioral RTs, as well as between neural response amplitude and both behavioral RTs and accuracy. These results suggest that the human V1, with neurons tuned to various features, generates saliency signals during the early stages of feedforward processing. These saliency signals are associated with the perceptual attraction of attention by feature pop-out. Importantly, these effects remain consistent even when the feature is unlikely to be resolved by SC neurons, thus suggesting that the V1SH holds true over alternative hypotheses.

To avoid circular analysis between electrode selection and ERP analysis, we selected the electrodes in an independent localizer session (Button, 2019). The C1 in the localizer session did not guarantee an effect of feature singleton in the main EEG session. Therefore, the singleton-specific effect could be rigorously tested. The polarity reversal characteristic results from the anatomical features of the calcarine sulcus. Together with a very short peak/trough time of the effect (70/76 ms), the topographic maps and source estimation results, the evidence strongly suggests a sensory-driven bottom-up process in V1 (Di Russo *et al*., 2002; Rauss *et al*., 2011b).

The polarity reversal as a unique characteristic of EEG signals from V1 is questioned in several papers. Particularly, a simulation study argues that ventral and dorsal V2/V3 sources can generate reversed polarity as well (Ales *et al*., 2010). However, subsequent studies suggest that simulated V1 sources produce negative activity for UVF stimuli and positive activity for LVF stimuli, consistent with the empirical data. Simulated V2/V3 sources, on the other hand, produce the opposite correspondence (Kelly *et al*., 2013b; Slotnick, 2018). Therefore, since the singleton-specific effects match the topographic activation profile of V1 source rather than that of V2/V3 source, they should primarily reflect V1 activity. In the main EEG session, we observed a negative C1 in the general ERPs (not the singleton-specific effects) for both the UVF and LVF singleton conditions (Figures 2D and 2E). This result was most likely elicited by the full-screen stimulation used. Because the LVF representation extends into the lower bank of the calcarine sulcus in many individuals, full-screen or horizontal-meridian stimulation (Foxe *et al*., 2008) causes dominant neural activity from the lower bank of the sulcus and results in negative voltages in occipital electrodes (Kelly *et al*., 2013a; Rauss *et al*., 2011b).

Our control analysis shows that the neural effect of saliency from the eye of origin cannot be explained by possible physical or psychophysical differences in luminance between the two eyes. With training, subjects may learn to discriminate the eye of origin based on subtle yet detectable binocular differences. Subjects often report brighter images in the dominant eye than in the non-dominant eye (Porac and Coren, 1984). Our analysis indicates that this potential binocular difference cannot generate a systematic early occipital effect comparable to the observed saliency effect. In fact, attention captured by eye-of-origin singletons retains even when luminance was randomly jittered across all bars to exclude any possible contribution of luminance difference to saliency (Zhaoping, 2008).

### V1 and exogenous attention

Behavioral studies have demonstrated the saliency effect based on eye-of-origin features (Zhaoping, 2008, 2012, 2018). Our experiment repeated this effect and further provided direct evidence of its neural substrate and response dynamics, suggesting a critical role of V1 feedforward mechanism. The neural response effects observed in our study reflect surround modulation mechanisms in visual saliency (Desimone and Duncan, 1995; Li, 1999) rather than the mere physical presence of an object or feature. Similar to many other studies (Chen *et al*., 2020; Liu *et al*., 2025; White *et al*., 2017; Yan *et al*., 2018; Zhang *et al*., 2012b), this is demonstrated by comparing the singleton-presence and baseline uniform conditions, which isolate the impact of center-surround feature differences.

Our results contradict previous proposals of other cortical areas, including IPS, FEF and V4, as sources of saliency signals, considering their longer response latency, different EEG activation topography, and lack of monocular neurons. Other studies have provided converging evidence supporting this conclusion. For example, the IPS typically exhibit attentional modulations with a latency over 100 ms (Gottlieb *et al*., 1998). Although the parietal cortices play a causal role in regulating saliency-related behaviors (Chen *et al*., 2020), fMRI laminar analysis revealed that the saliency signals in IPS are transmitted from V1 superficial layers (Liu *et al*., 2025). Lesions in V4 impair the selection of nonsalient locations in a scene while sparing the detection of salient objects (Schiller and Lee, 1991). Similarly, damage to the frontal eye fields (FEF) disrupts voluntary visual tracking (Lynch, 1987) but minimally affects stimulus-driven saccadic orientation to salient locations (Schiller *et al*., 1987).

The results also contradict the hypothesis that the primate SC is necessary for saliency representation in V1. Electrophysiological recordings in monkeys have revealed earlier salience coding in the SC, with an onset latency of 65 ms compared to an onset latency of 139 ms in V1, suggesting that the saliency computation first occurs in the SC with its surround suppression mechanisms and generates feedback signals to V1 (White *et al*., 2017). Since it is difficult to record SC responses with high temporal resolutions in humans, we focused on V1 responses to eye-of-origin singletons, which are unlikely to be resolved by SC neurons (Finlay *et al*., 1976a; Schiller *et al*., 1974; Zhaoping, 2008). Our results demonstrate that this eye-of-origin feature still generates an early attention-related V1 effect in the first-response component of the occipital cortex. The latency (onset latency: 58–70 ms, peak latency: 70-78 ms) and amplitude are not significantly different from those associated with saliency from orientation features recognizable by SC neurons (Baumann *et al*., 2023; Chen and Hafed, 2018). This latency is similar to that seen in previous experiments with orientation singletons in humans (60-80 ms) and monkeys (40-80 ms) (Yan *et al*., 2018; Zhang *et al*., 2012b). Additionally, our behavioral data do not support temporal or upper visual field biases, which have been demonstrated in neuronal activity of primate SCs (Hafed and Chen, 2016; Sylvester *et al*., 2007). Thus, these findings suggest that saliency representation in V1 relies on its feedforward intracortical mechanisms instead of SC involvement.

The discrepancy between our results and those of White *et al*. (2017) regarding the latency of the earliest V1 saliency effect likely stems from the difference in what each method measures. A computational model suggests that saliency computation involves localized neuronal subgroups selective to pop-out features and feature-specific lateral inhibition from a large group of nearby, feature-selective neurons and inhibitory interneurons (Li, 1999; Zhaoping and Zhe, 2015). EEG may reflect large-scale, synchronous activity from all these neural populations and intracortical processing (Cohen, 2017). In contrast, White *et al*. (2017) recorded the spiking activity of V1 neurons with a response field exactly at the singleton location. We speculate that pooling neurons with diverse feature preferences and ignoring nearby neurons and interneurons may have averaged out the small, prominent early saliency signals present only in neurons selective to the pop-out feature. As with V1 neurons grouped by their feature preference, a clear saliency signal emerges within the initial response window (Yan *et al*., 2018).

Consistent with the feature-integration effect in visual saliency in the present and previous behavioral tests (Treisman and Gelade, 1980; Zhaoping, 2008), we observed an enhanced neuronal response amplitude in the EO conditions compared with the E and O conditions. Supporting a proposed model of saliency calculation derived from V1SH (Koene and Zhaoping, 2007; Zhaoping and Zhe, 2015), our results suggest that neuronal response facilitation by feature conjunction occurs at V1. This facilitation may occur through a race between two groups of neurons tuned to different features and/or neurons tuned to both features conjunctively (i.e., monocular oriented neurons). In contrast, we did not observe a systematic shortening of neural latency in the EO conditions, suggesting a similar timing for saliency calculation in neural populations tuned to different features. Similarly, previous results show that the latency of the earliest saliency signals does not shift with increased feature contrasts or task practice (Yan *et al*., 2018; Zhang *et al*., 2012b). Note that this saliency signal should be differentiated from the attentional modulation effect, which occurs later and can exhibit shifted latency related to attentional strength (Galashan *et al*., 2013; Yan *et al*., 2018).

In monkey V1, neuronal responses are facilitated by exogenous cues approximately 80–180 ms after probe onset (Wang *et al*., 2015). A similar effect occurs 80–200 ms after cue onset when the target is the cue itself, following the saliency calculation response (Yan *et al*., 2018). Transcranial magnetic stimulation (TMS) at V1/V2 during probe presentation eliminates the effects of exogenous attention without altering endogenous attention modulation. In contrast, TMS at the FEF weakens endogenous attention (Fernández and Carrasco, 2020; Fernández *et al*., 2023). These findings suggest that V1 plays a causal and immediate role in mediating exogenous attention but is not critical for endogenous attention until feedback from higher visual areas arrives. Our results provide a possible explanation for this dissociation between exogenous and endogenous attention: the feedforward saliency signals in V1 may exert their effect directly and most efficiently within V1, well before the endogenous signals transmitted through intercortical connections. Our results also support a bottom-up mechanism for eye-specific attention in V1 (Zhang *et al*., 2012a). With a similar intracortical mechanism suggested by the present study, eye-specific attention can be generated and represented within V1 monocular neurons, resulting in a shift in ocular dominance (Song *et al*., 2023; Wong *et al*., 2021; Wong *et al*., 2025).

### Correlation between neural responses and behaviors

In the EEG experiment, we used an orthogonal fixation task to isolate the early saliency signals from potential top-down effects and eye-movements noise as much as possible (Rauss *et al*., 2011a; Slotnick, 2018; Wolf *et al*., 2021). The observed effects were similar between the E and O conditions despite the difference in visibility between eye-of-origin and orientation singletons, suggesting the dominance of bottom-up signal. This finding supports the notion that bottom-up attention occurs during a pre-attentive phase (Liang *et al*., 2023; Wolfe, 2021; Zhang *et al*., 2012b; Zhaoping and Guyader, 2007). In this phase, the eye of origin is a basic feature that guides attention, and its effect is comparable to that of other visible features, such as color and orientation (Zhaoping, 2008, 2012).

On the other hand, we used a detection task similar to many previous studies (Wolfe, 2021; Yan *et al*., 2018; Zhaoping, 2008; Zhaoping and Zhe, 2015) to reflect attention capture in the behavioral experiment. The performance was partially affected by high-level perception. Particularly, the low performance in the E condition was caused by the difficulty identifying rather than “finding” the eye-of-origin singletons (Zhaoping, 2012). Despite different tasks in the EEG and behavioral experiments, studies show that saliency signals occur in V1 initial responses regardless of the detection task, task familiarity, or access to awareness. Instead, a later component that appears during the 80-200 ms period is relevant to these factors (Knierim and van Essen, 1992; Yan *et al*., 2018; Zhang *et al*., 2012b). Therefore, we predict largely shared saliency processing in the two experiments. This is later confirmed by our correlation results.

We found that the amplitude of the early occipital effect was positively correlated with accuracy and negatively correlated with RTs in the detection task. Similarly, the cueing effect of orientation singletons has been found to correlate with V1 activity across observers (Liu *et al*., 2025; Zhang *et al*., 2012b). Additionally, correlations have been observed between C1 amplitude and attention-related contrast appearance (Pan *et al*., 2024). These results suggest that the amplitude of V1 activity drives the strength of bottom-up attentional attraction. Furthermore, our results showed that the speed-adjusted accuracy had the strongest correlation with the neural response. Although this is an oversimplified integration of behavioral accuracy and RTs (Liesefeld and Janczyk, 2019), it implies that V1 saliency strength contributes to behavioral efficiency, which may manifest as either correctness or RT, depending on the speed-accuracy tradeoff.

Interestingly, we also observed that the latency of singleton-specific occipital activity was correlated with behavioral RTs but not accuracy, implying that neuronal latency specifically contributes to behavioral speed. Similarly, neuronal latency in monkey MT is correlated with behavioral RT when detecting a speed change. This correlation can be explained by the variability of neuronal responses prior to the speed change, indicating that changes in network noise cause variations in latency (Galashan *et al*., 2013). Trial-by-trial correlations have also been found between FEF neural latency and behavioral latency in smooth pursuit. Preparatory activity at baseline substantially influences these latencies (Lee *et al*., 2019). Considering these findings, the between-subject correlations observed in our experiment may reflect individual variations in baseline neural network activity. This baseline state in V1 partially determines the speed of saliency generation, which then affects the speed of saliency-dependent behaviors, though it is not related to enhanced saliency signals caused by increased (Yan *et al*., 2018; Zhang *et al*., 2012b) or combined feature contrasts.

### Limitation

The SC was not directly recorded in the present study, although its responses were presumably manipulated by stimulus conditions based on prior physiological knowledge. Therefore, while our conclusion that V1 performs feedforward saliency processing is very likely correct considering the short latency, we cannot completely rule out possibilities such as simultaneous saliency processing in the SC. Future studies with invasive recordings on human patients might be necessary to better understand these subcortical mechanisms. The head-mounted display limited our ability to record eye movements. Nevertheless, analysis of EOG/EEG responses showed that the observed effect was unlikely caused by systematic artifacts of eye movement. Studies have shown that the C1 component is sensitive to slight eye deviations at stimulus onset (Wolf *et al*., 2021). In our study, there should be no such systematic eye deviations because the subjects could not know the singleton position until trial start.

## Materials and Methods

### Participants

Thirty-one healthy right-handed subjects (15 females, 16 males, aged 18–35 years, mean age = 26.4 ± 4.2 years) with normal or corrected-to-normal vision were initially recruited. Written informed consent was obtained from all participants prior to their participation. All participants were naive to the experimental hypotheses and had no history of neurological or visual disorders. Two subjects failed to complete the EEG data acquisition due to reported dizziness and frequent baseline drift. The study protocol was approved by the Human Subjects Review Board of Changsha Aier Eye Hospital (Approval Number: (2023) KYPJ027) and was conducted in accordance with the ethical standards outlined in the Declaration of Helsinki.

### Apparatus

Visual stimuli were generated with Psychtoolbox (Brainard, 1997) in MATLAB (MathWorks, USA) and presented on a head-mounted display (Goovis G3 Max, China, 60 Hz, 2560 × 1440 pixels for each screen/eye) with a simulated viewing distance of 20 meters. The head position was stabilized using a chin rest. Screens were calibrated with a spectroradiometer (Spectraval 1511, JETI Technische Instrumente GmbH, Germany). EEG (and EOG) data were acquired using an EGI GES-400 EEG system with a 64-channel HydroCel GSN 130 saline-based electrode cap (Magstim Inc, USA). The impedance of all parietal and occipital electrodes, which were the focus of our study, was adjusted and maintained below 50 kΩ (preferentially below 30 kΩ) throughout the recordings, while the impedance of other electrodes was maintained below 100 kΩ. Neural signals were amplified with a gain of 500K and digitized at a sampling rate of 500 Hz. Online recordings utilized a vertex reference (Cz). The delay of actual stimuli presentation was measured using an audio/visual (AV) device (Magstim Inc, USA) toolkit.

### Stimuli and procedure

The visual stimuli consisted of texture patterns comprising a matrix of 23 × 31 white bars (10 cd/m²) presented against a dark background (1 cd/m²). The entire stimulus array subtended a visual angle of 36.8° × 49.6°. Each bar was a rectangle of 0.2° × 1.1° in visual angle, separated by a center-to-center distance of 1.6°, with random jitters of 0.1° at maximum in both horizontal and vertical directions. The eye of origin of a bar was defined as the luminance contrast between the inputs of the left and right eyes:

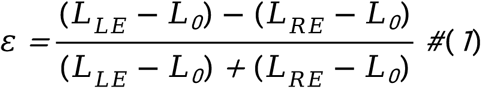

where *L_LE_* and *L_RE_* denote the luminance of the bar in the left eye and right eye, respectively, and *L_0_* denotes the luminance of screen background. In each trial, *ε* was randomly set as 0.7 or −0.7, orientation was randomly set between 0° to 180°. All bars, except for one singleton, shared the same *ε* and orientation. The singleton bar differed in terms of *ε* (E), orientation (O), both *ε* and orientation (EO), or absent (U). In O and EO conditions, the singleton bar was tilted ±36° from the background bars. It was presented at one of four locations on the diagonal lines, with an eccentricity of 9°. Between the bar stimulus, small dots were binocularly presented to help in binocular alignment.

All subjects completed a behavioral visual detection test before the EEG experiment. In this test, each trial commenced with a fixation period that randomly varied between 1200 ms and 1400 ms, followed by a stimulus presentation of 50 ms. Participants were required to maintain central fixation and press left/right buttons to report when a singleton appeared on the left/right half of the screen. Specific instructions were given to the participants in addition to the brief stimulus to ensure that the responses were reflexive without special search strategies. Each subject completed 64 trials for each of E, O, and EO conditions, with two practice blocks (total of 192 trials) before the formal experiment. The singleton feature condition and the singleton location, the *ε* value of the background bars, and sign of orientation contrast between the singleton and background bars were pseudorandomized in the trial sequence.

We employed similar visual stimuli in the EEG experiment as those in the behavioral experiment. In the main session, each trial began with a central fixation for a randomized duration between 800 ms and 950 ms, followed by a 50-ms stimulus and a subsequent 300-ms fixation-only period. Subjects were required to fixate their gaze at the center of the screen and identify an occasional color change of fixation. The color change randomly occurred for 200 ms in every 5–20 trials. The program proceeded to the next trial when a button press was registered or a maximum time of 1800 ms was reached. This task was set to maintain the eye gaze and monitor the attention of the observer. The main session included 576 trials for each of the U, E, O and EO conditions, respectively. Before the main sessions, a localizer session was performed to select the electrodes of interest. In this session, a single star-shaped object composed of four bars was presented at one of four locations, which were the same as the singleton locations in the main session. The localizer session consisted of 768 stimulus-present trials and 256 stimulus-absent trials.

Before the experiment, each of the two screens in the head-mounted display were calibrated at all singleton positions. This was done to ensure precise luminance display, thus avoiding potential differences in binocular luminance between the singleton and background bars (particularly around the singleton positions) in the E and EO conditions (i.e., when the singleton and background bars were presented in different screens/eyes). This was critical to the measurement of saliency by eye-of-origin contrasts, as there could be confounding effect of luminance contrasts due to different output luminance between screens if uncalibrated. Although we only performed the calibration at the singleton location, the nonuniformity (existing in any screen) at other locations was small and spatially smooth; thus, it induced little effect on saliency. Further, the screens were preheated for at least one hour both before the calibration and the experiment.

### EEG data preprocessing and electrode selection

EEG data was preprocessed with EEGLAB (Delorme and Makeig, 2004) and included the following steps: The raw data was filtered using a bandpass noncausal filter of 0.01 Hz–40 Hz and re-referenced to the average of bilateral mastoids. The cut-off value of 0.01 Hz was selected to minimize potential bias caused by the high-pass filter, especially with regard to the early ERP components (Acunzo *et al*., 2012). Independent component analysis (ICA) (runica.m) was utilized to detect and remove blink and eye-movement artifacts (probability >90%, recognized by ICLable). Continuous data was then segmented into epochs, spanning from 100 ms before to 250 ms after the stimulus onset. Baseline correction was performed using the mean voltage during the 100-ms pre-stimulus interval. Trials with data exceeding ±50 μV, large drifts or large baseline noise (baseline data drift exceeding 25 μV within 20 ms; baseline data range exceeding the mean plus three standard deviations of the baseline data range across all trials) were rejected. After artifact rejection, there remained 87.5%-99.7% trials in the U condition, 89.4%-99.7% trials in the E condition, 87.8%-99% in the O condition, and 88.4%-98.8% trials in the EO condition.

The occipital electrodes of interest were selected to best capture the earliest visual responses. This was done by selecting the electrodes with the strongest C1 amplitudes in the independent localizer session, as C1 was a characteristic visual evoked potential that represented the earliest V1 responses (Di Russo *et al*., 2002; Rauss *et al*., 2011b). With strong prior knowledge of C1 and to avoid the possible artifacts in frontal electrodes caused by the head-mounted display, we restricted the electrode selection to the parieto–occipital region. Based on the remarkable polarity reversal feature and typical time range, we first identified C1 in each electrode as the largest peak or trough during 20–100 ms elicited by the stimuli in the lower visual field (LVF) or upper visual field (UVF). Then, the difference between the peak and trough was obtained as the C1 strength for a particular electrode. Finally, six posterior electrodes that demonstrated the largest C1—E33 (PO3), E39 (O2), E34 (PZ), E36 (POZ), E37 (OZ), and E38 (PO4)—were selected and utilized in the analysis of the main EEG session.

VEOG was recorded as the differential voltage between two electrodes placed above the eyes and two electrodes placed below the eyes in the current system (Ai *et al*., 2016). We obtained the event-related VEOG responses using the same processing steps as for the occipital electrodes, but without ICA analysis to retain the ocular components in the data. The results were analyzed for singletons in the UVF and LVF separately. To examine the potential effect of vertical eye movements on the reference electrodes, we also checked the responses in the mastoid electrodes without re-referencing and ICA analysis.

We measured the delay of actual stimuli presentation from the received trigger in EEG recordings in a separate test before the experiment using the same stimuli program. The luminance change at stimulus onset was detected with a photosensor which sent a TTL pulse to the EEG amplifier. Then, the mean time difference (14 ms, range: 6-23 ms in 336 trials) between this TTL pulse and its corresponding trigger sent from the stimulus computer was calculated and compensated for in the data analysis stage (i.e., the actual stimulus onset time was corrected as the registered time plus 14 ms). We confirmed that the trial-by-trial variation in delay time was caused by the head-mounted display. Despite this variation, our conclusions based on average responses across trials were unaffected.

### Singelton-specific effect in individual data

We searched (MATLAB findpeaks function) for the largest effect peak (singletons in the LVF) or trough (singletons in the UVF) within the time window 20–100 ms for each subject separately. The peak/trough amplitude was determined as the average amplitude over the peak/trough point along with four neighboring time points. Then the effect combining the UVF and LVF was acquired as the difference between the peak and trough. When only one of the peak and trough could be detected properly (3-5 subjects in different singleton conditions), the combined effect value was set to the detected one. To determine if the effect was random noise, we performed a control analysis in which we searched for the peak/trough in the EEG data with a reversed polarity. This was equivalent to searching for troughs when singletons were present in the LVF and searching for peaks when singletons were present in the UVF. If the detected effect values were random noise, no systematic difference in amplitude would be expected between the original and reversed-polarity data.

### Source estimation

We utilized the DIPFIT toolbox of EEGLAB for dipole source estimation. A head model was first constructed using publicly available average brain images from the Montreal Neurological Institute (MNI) database, which consisted of a high-quality anatomical MRI of a single representative subject with a voxel size of 1 × 1 × 1 mm. The co-registration of the head model and electrode locations was completed based on fiducial and Cz points with a semi-automatic procedure. Then, we constructed the equivalent current dipole model using the DIPFIT inverse function. This function first scanned the three-dimensional grid of the head model to determine the acceptable starting positions, and then utilized a nonlinear optimization algorithm to fit the exact dipole position. To increase the signal-to-noise ratio (SNR) and considering similar responses across different singleton conditions, we obtained the topographic maps and performed source localization using the averaged signals from the E, O, and EO conditions at the peak latencies of the attention effect — 70 ms for the upper visual field and 76 ms for the lower visual field. We utilized symmetrical dipole models assuming that similar responses would be obtained for two hemispheres in the early stages of visual processing.

### Statistics

To detect the early attentional effect caused by the feature singletons, we tested the difference between the singleton-present conditions (E, O, EO) and singleton-absent condition (U) using cluster-based permutation methods on the average ERP responses within a time interval ranging from 20 ms to 100 ms (Figures 2D and 2G). In each of a total of 5000 permutations, we randomly flipped the signs of each participant’s data to simulate the random effects under the null hypothesis. The t-test was performed at each time point of the permuted data to identify consecutive clusters of time points that showed significant differences (time point p < 0.05). Then, the null distribution was generated from the length of the largest cluster in each permutation. Finally, we calculated the length of the cluster using the non-permuted original data, with its p-value obtained as the position of the cluster length in the null distribution. This method corrected the multiple comparison problem caused by simultaneous tests at different time points. In addition, we further adjusted the statistical threshold using Bonferroni-Holm method for multiple comparisons in the six different conditions. The beginning time of the significant cluster was recognized as the onset latency of the attentional effect.

Another permutation test was performed on the singleton-specific effect detected in individual subject data: in each of a total 5000 permutations, the polarity of the individual data was randomly reversed. Then, the largest effect peak/trough was searched in each permutation, and the mean amplitude across subjects was calculated. Subsequently, the null distribution was established from these mean amplitudes in all permutations. Finally, the mean amplitudes calculated using the original data were compared with the null distribution, and the p-value was obtained as its position in this distribution. The statistical threshold was adjusted using the Holm method for multiple comparisons in the three singleton conditions.

ANOVA and power analysis were performed using JASP (Version 0.19.3; JASP Team (2024)). Post-hoc pairwise comparisons were corrected for multiple testing using the Holm method. Unless otherwise stated, all reported p-values in the Results section were corrected for multiple comparisons.

The sample size was determined based on a preliminary experiment that resulted in an effect size of d = 0.55–1.03 for the singleton-specific effects revealed by comparing the original and polarity-reversed EEG data (similar as Figure 3). With an alpha level of 0.05 and a desired power of 0.8, the power analysis indicated a minimum sample size of 28. Given the final sample size of 29 effective participants, our data were sensitive enough to detect an effect size of d = 0.539 or larger.

## Code and data accessibility

The data and analysis code have been archived on the Open Science Framework (OSF). The registration will be made publicly available upon publication.

## Author Contributions

P.Zhang, Z.Yang and J.Zou designed research; C.Wu, H.Li, X.Wang and Z.Yin performed research; C.Wu, X.Li, Z.Wang and J.Zou analyzed data; X.Li, P.Zhang, Z.Yang and J.Zou edited the paper; C.Wu and J.Zou wrote the paper.

## Acknowledgment

This research was funded by Provincial Natural Science Foundation of Hunan (Grant No.2024JJ9001), Hunan Xiangjiang Philanthropy Foundation (Grant No.KY24016) and Clinic Research Foundation of Aier Eye Hospital Group (Grant No.AIM2301D04).

## Supplementary materials

**Figure S1.**
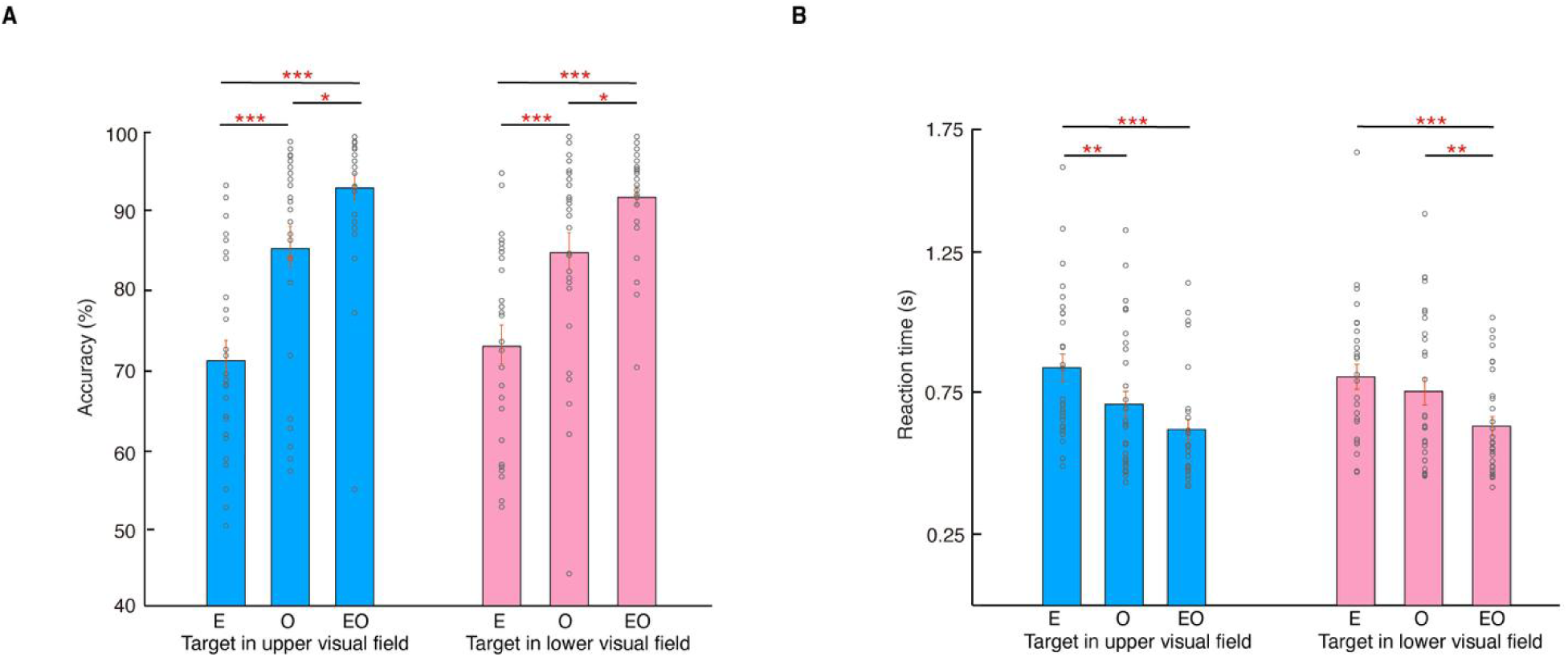
Detection accuracy (A) and reaction time (B) in the upper and lower visual fields. Gray circles indicate data of individual subjects. Error bars represent SEM. *, **, and *** indicate Holm-corrected p < 0.05, p < 0.01, and p < 0.001.

**Figure S2.**
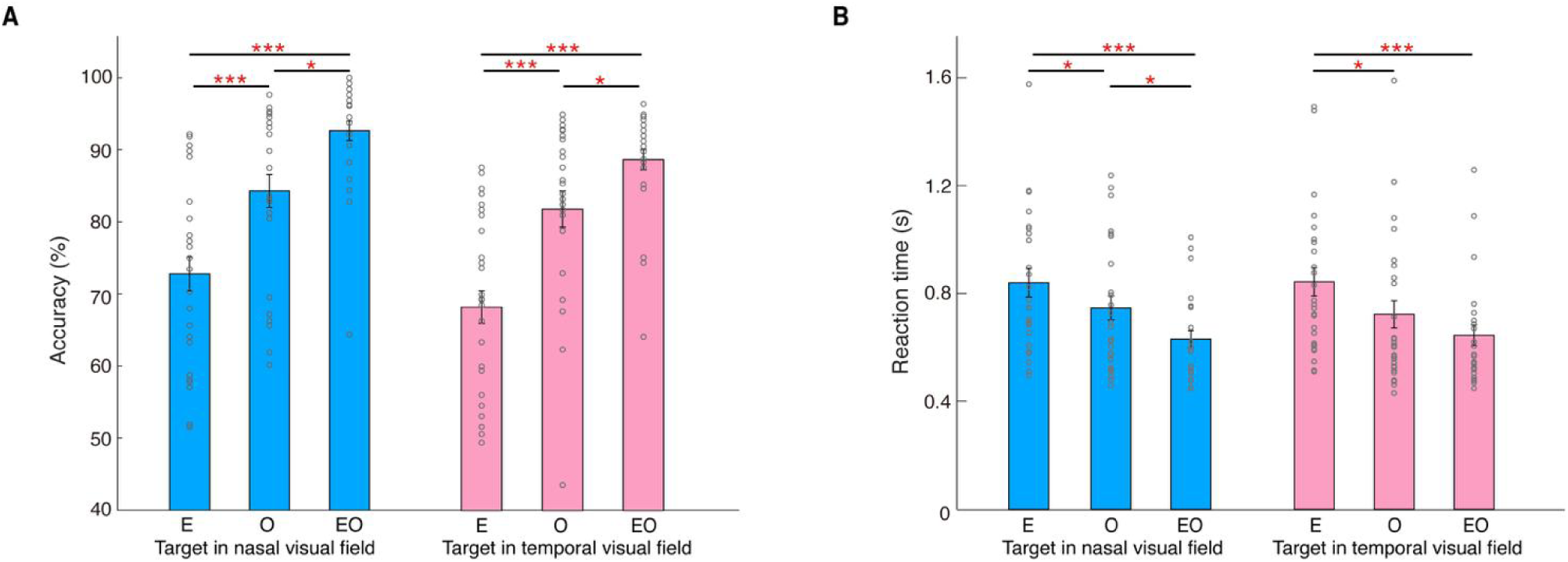
Detection accuracy (A) and reaction time (B) in the nasal and temporal visual fields. Gray circles indicate data of individual subjects. Error bars represent SEM. * and *** indicate Holm-corrected p < 0.05 or p < 0.001.

**Figure S3.**
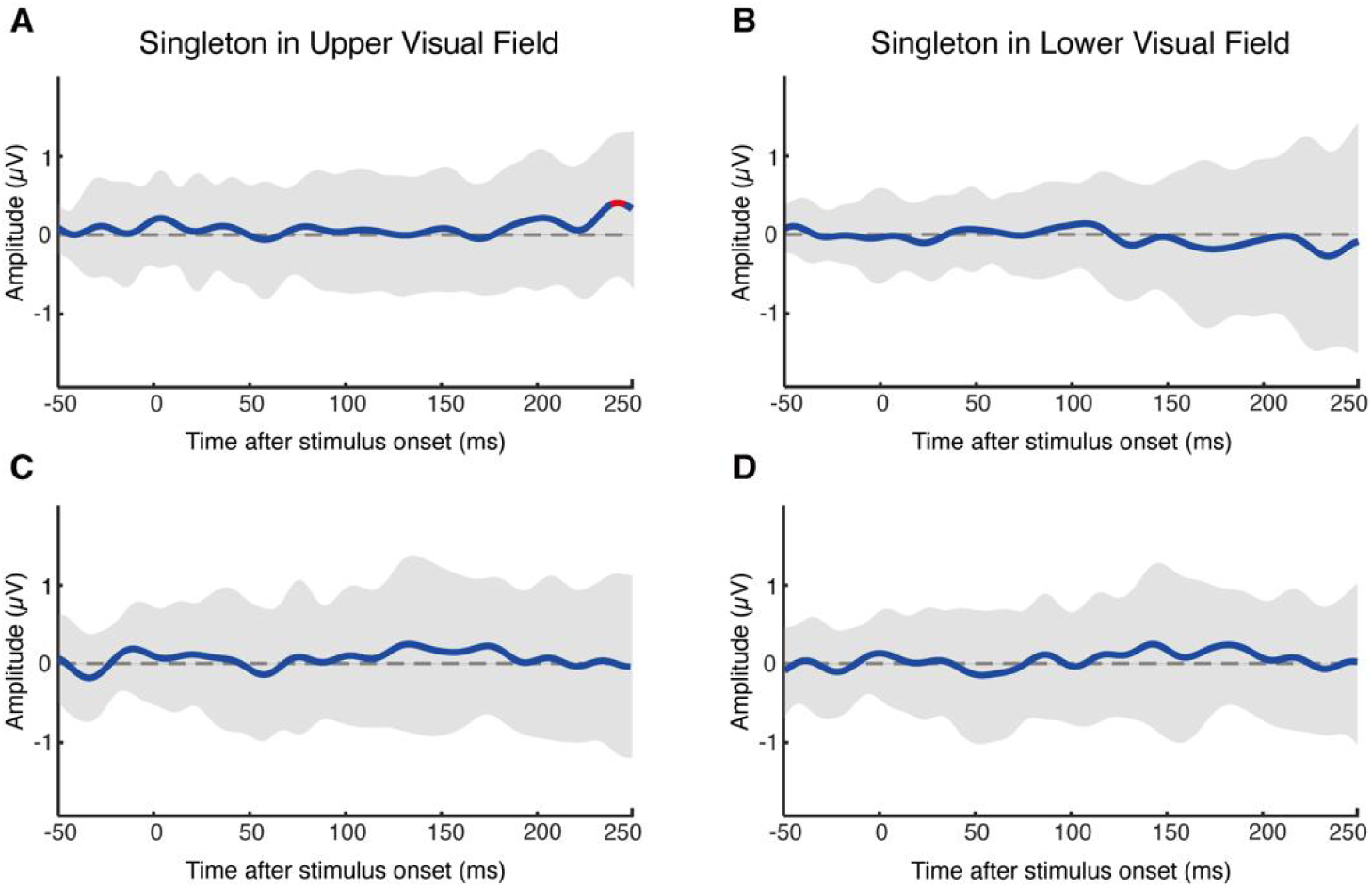
Eye movement artifacts in EEG data. **(A, B)** Vertical electrooculogram (VEOG). VEOG was derived from the differential voltage recorded between electrodes placed above and below the eyes. **(C, D)** Responses in the mastoid electrodes. Singleton-specific responses (singleton-presence conditions vs. singleton-absence conditions) were obtained for singletons in the upper (A, C) and lower (B, D) visual fields, respectively. Gray shaded areas represent SEM. Red dots indicate uncorrected p < 0.05.

**Figure S4.**
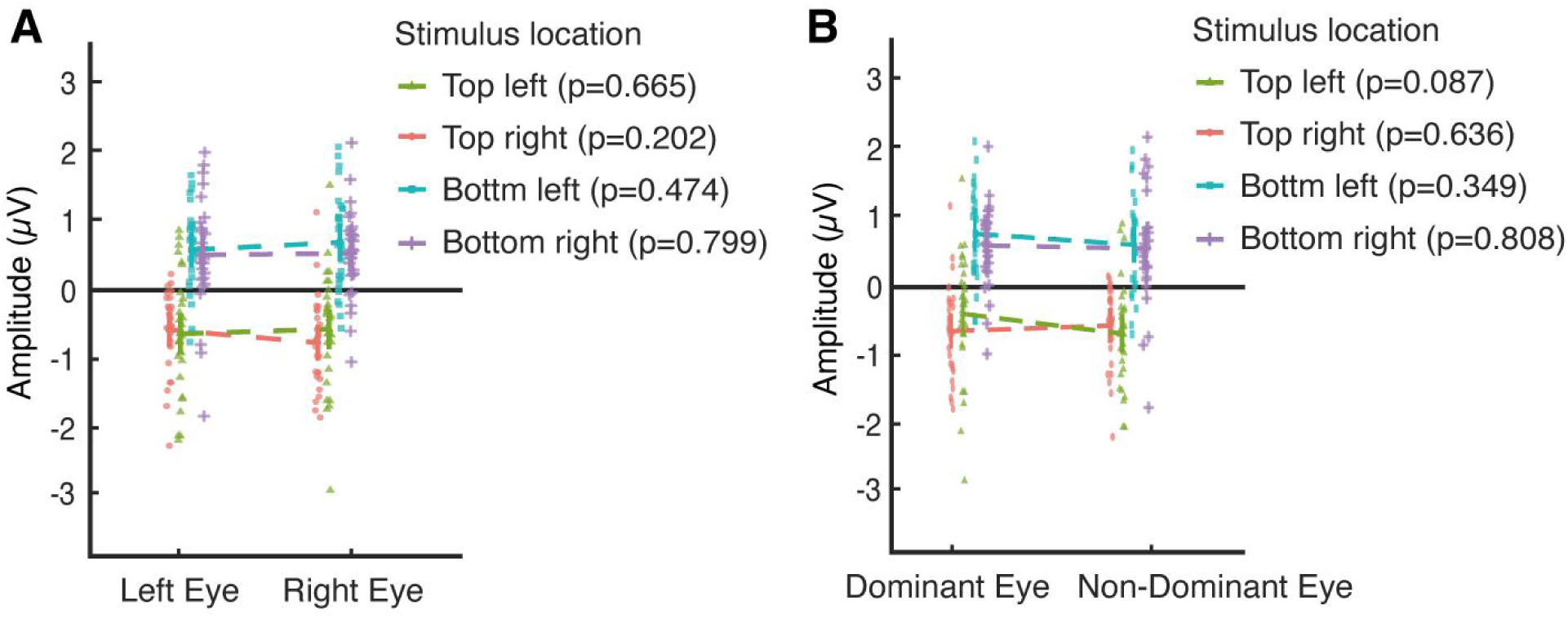
C1 amplitudes elicited by left-eye and right-eye stimuli in the localizer session. The data were grouped by either **(A)** left and right eye, or **(B)** dominant and non-dominant eye. Uncorrected p-values were provided for the comparison at each stimulus location (the same as the singleton location in the main EEG session). Error bars represent SEM, with individual data points overlaid.

## References

Acunzo, D.J., MacKenzie, G., van Rossum, M.C.W., 2012. Systematic biases in early ERP and ERF components as a result of high-pass filtering. J. Neurosci. Methods 209, 212–218.

Ai, G., Sato, N., Singh, B., Wagatsuma, H., 2016. Direction and viewing area-sensitive influence of EOG artifacts revealed in the EEG topographic pattern analysis. Cogn. Neurodyn. 10, 301–314.

Ales, J.M., Yates, J.L., Norcia, A.M., 2010. V1 is not uniquely identified by polarity reversals of responses to upper and lower visual field stimuli. Neuroimage 52, 1401–1409.

Baumann, M.P., Bogadhi, A.R., Denninger, A.F., Hafed, Z.M., 2023. Sensory tuning in neuronal movement commands. Proc Natl Acad Sci U S A 120, e2305759120.

Button, K.S., 2019. Double-dipping revisited. Nat. Neurosci. 22, 688–690.

Chen, C.Y., Hafed, Z.M., 2018. Orientation and Contrast Tuning Properties and Temporal Flicker Fusion Characteristics of Primate Superior Colliculus Neurons. Front Neural Circuits 12, 58.

Chen, X., Zirnsak, M., Vega, G.M., Govil, E., Lomber, S.G., Moore, T., 2020. Parietal Cortex Regulates Visual Salience and Salience-Driven Behavior. Neuron 106, 177–187 e174.

Cohen, M.X., 2017. Where Does EEG Come From and What Does It Mean? Trends Neurosci. 40, 208–218.

Corbetta, M., Shulman, G.L., 2002. Control of goal-directed and stimulus-driven attention in the brain. Nat Rev Neurosci 3, 201–215.

Delorme, A., Makeig, S., 2004. EEGLAB: an open source toolbox for analysis of single-trial EEG dynamics including independent component analysis. J Neurosci Methods 134, 9–21.

Desimone, R., Duncan, J., 1995. Neural Mechanisms of Selective Visual Attention. Annu. Rev. Neurosci. 18, 193–222.

Di Russo, F., Martínez, A., Sereno, M.I., Pitzalis, S., Hillyard, S.A., 2002. Cortical sources of the early components of the visual evoked potential. Hum. Brain Mapp. 15, 95–111.

Dougherty, K., Cox, M.A., Westerberg, J.A., Maier, A., 2019. Binocular Modulation of Monocular V1 Neurons. Curr. Biol. 29, 381–391 e384.

Fernández, A., Carrasco, M., 2020. Extinguishing Exogenous Attention via Transcranial Magnetic Stimulation. Curr. Biol. 30, 4078–4084.e4073.

Fernández, A., Hanning, N.M., Carrasco, M., 2023. Transcranial magnetic stimulation to frontal but not occipital cortex disrupts endogenous attention. Proceedings of the National Academy of Sciences 120, e2219635120.

Finlay, B.L., Schiller, P.H., Volman, S.F., 1976a. Quantitative studies of single-cell properties in monkey striate cortex. IV. Corticotectal cells. J. Neurophysiol. 39, 1352–1361.

Finlay, B.L., Schiller, P.H., Volman, S.F., 1976b. Quantitative studies of single-cell properties in monkey striate cortex. IV. Corticotectal cells. J Neurophysiol 39, 1352–1361.

Foxe, J.J., Strugstad, E.C., Sehatpour, P., Molholm, S., Pasieka, W., Schroeder, C.E., McCourt, M.E., 2008. Parvocellular and Magnocellular Contributions to the Initial Generators of the Visual Evoked Potential: High-Density Electrical Mapping of the “C1” Component. Brain Topogr. 21, 11–21.

Galashan, F.O., Saßen, Hanna C., Kreiter, Andreas K., Wegener, D., 2013. Monkey Area MT Latencies to Speed Changes Depend on Attention and Correlate with Behavioral Reaction Times. Neuron 78, 740–750.

Gottlieb, J.P., Kusunoki, M., Goldberg, M.E., 1998. The representation of visual salience in monkey parietal cortex. Nature 391, 481–484.

Hafed, Z.M., Chen, C.Y., 2016. Sharper, Stronger, Faster Upper Visual Field Representation in Primate Superior Colliculus. Curr. Biol. 26, 1647–1658.

Itti, L., Koch, C., 2001. Computational modelling of visual attention. Nat Rev Neurosci 2, 194–203.

Kelly, S.P., Schroeder, C.E., Lalor, E.C., 2013a. What does polarity inversion of extrastriate activity tell us about striate contributions to the early VEP? A comment on Ales et al. (2010). Neuroimage 76, 442–445.

Kelly, S.P., Vanegas, M.I., Schroeder, C.E., Lalor, E.C., 2013b. The cruciform model of striate generation of the early VEP, re-illustrated, not revoked: A reply to Ales et al. (2013). Neuroimage 82, 154–159.

Knierim, J.J., van Essen, D.C., 1992. Neuronal responses to static texture patterns in area V1 of the alert macaque monkey. J. Neurophysiol. 67, 961–980.

Koene, A.R., Zhaoping, L., 2007. Feature-specific interactions in salience from combined feature contrasts: evidence for a bottom-up saliency map in V1. J. Vis. 7, 6 1–14.

Krummenacher, J., Muller, H.J., Heller, D., 2001. Visual search for dimensionally redundant pop-out targets: evidence for parallel-coactive processing of dimensions. Percept Psychophys 63, 901–917.

Lee, J., Darlington, T.R., Lisberger, S.G., 2019. The Neural Basis for Response Latency in a Sensory-Motor Behavior. Cereb. Cortex 30, 3055–3073.

Li, Z., 1999. Contextual influences in V1 as a basis for pop out and asymmetry in visual search. Proc. Natl. Acad. Sci. U. S. A. 96, 10530–10535.

Liang, J., Maher, S., Zhaoping, L., 2023. Eye movement evidence for the V1 Saliency Hypothesis and the Central-peripheral Dichotomy theory in an anomalous visual search task. Vision Res. 212, 108308.

Liesefeld, H.R., Janczyk, M., 2019. Combining speed and accuracy to control for speed-accuracy trade-offs(?). Behavior Research Methods 51, 40–60.

Liu, C., Liu, C., Huber, L., Zhaoping, L., Zhang, P., 2025. The superficial layers of the primary visual cortex create a saliency map that feeds forward to the parietal cortex. PLoS Biol. 23, e3003159.

Lynch, J.C., 1987. Frontal eye field lesions in monkeys disrupt visual pursuit. Exp Brain Res 68, 437–441.

Nothdurft, H., 2000. Salience from feature contrast: additivity across dimensions. Vision Res. 40, 1183–1201.

Ono, H., Barbeito, R., 1985. Utrocular discrimination is not sufficient for utrocular identification. Vision Res. 25, 289–299.

Pan, W.N., Zhao, Y.W., Luo, Z.X., Chen, Y., Cai, Y.C., 2024. Attention modulates early visual processing: An association between subjective contrast perception and early C1 ERP component. Psychophysiology 61, e14507.

Plöchl, M., Ossandón, J.P., König, P., 2012. Combining EEG and eye tracking: identification, characterization, and correction of eye movement artifacts in electroencephalographic data. Front. Hum. Neurosci. Volume 6 - 2012.

Porac, C., Coren, S., 1984. Monocular asymmetries in vision: A phenomenal basis for eye signature. Canadian Journal of Psychology / Revue canadienne de psychologie 38, 610–624.

Rauss, K., Schwartz, S., Pourtois, G., 2011a. Top-down effects on early visual processing in humans: A predictive coding framework. Neurosci. Biobehav. Rev. 35, 1237–1253.

Rauss, K., Schwartz, S., Pourtois, G., 2011b. Top-down effects on early visual processing in humans: a predictive coding framework. Neurosci Biobehav Rev 35, 1237–1253.

Schiller, P.H., Lee, K., 1991. The role of the primate extrastriate area V4 in vision. Science 251, 1251–1253.

Schiller, P.H., Sandell, J.H., Maunsell, J.H., 1987. The effect of frontal eye field and superior colliculus lesions on saccadic latencies in the rhesus monkey. J Neurophysiol 57, 1033–1049.

Schiller, P.H., Stryker, M., Cynader, M., Berman, N., 1974. Response characteristics of single cells in the monkey superior colliculus following ablation or cooling of visual cortex. J. Neurophysiol. 37, 181–194.

Slotnick, S.D., 2018. The experimental parameters that affect attentional modulation of the ERP C1 component. Cogn. Neurosci. 9, 53–62.

Song, F., Lyu, L., Zhao, J., Bao, M., 2023. The role of eye-specific attention in ocular dominance plasticity. Cereb. Cortex 33, 983–996.

Sylvester, R., Josephs, O., Driver, J., Rees, G., 2007. Visual FMRI responses in human superior colliculus show a temporal-nasal asymmetry that is absent in lateral geniculate and visual cortex. J. Neurophysiol. 97, 1495–1502.

Treisman, A.M., Gelade, G., 1980. A feature-integration theory of attention. Cognitive Psychology 12, 97–136.

Veale, R., Hafed, Z.M., Yoshida, M., 2017. How is visual salience computed in the brain? Insights from behaviour, neurobiology and modelling. Philos Trans R Soc Lond B Biol Sci 372.

Wang, F., Chen, M., Yan, Y., Zhaoping, L., Li, W., 2015. Modulation of Neuronal Responses by Exogenous Attention in Macaque Primary Visual Cortex. J. Neurosci. 35, 13419–13429.

White, B.J., Kan, J.Y., Levy, R., Itti, L., Munoz, D.P., 2017. Superior colliculus encodes visual saliency before the primary visual cortex. Proc. Natl. Acad. Sci. U. S. A. 114, 9451–9456.

Wolf, M.-I., Bruchmann, M., Pourtois, G., Schindler, S., Straube, T., 2021. Top-Down Modulation of Early Visual Processing in V1: Dissociable Neurophysiological Effects of Spatial Attention, Attentional Load and Task-Relevance. Cereb. Cortex 32, 2112–2128.

Wolfe, J.M., 2021. Guided Search 6.0: An updated model of visual search. Psychon Bull Rev 28, 1060–1092.

Woltz, D.J., Was, C.A., 2006. Availability of related long-term memory during and after attention focus in working memory. Mem. Cognit. 34, 668–684.

Wong, S.P., Baldwin, A.S., Hess, R.F., Mullen, K.T., 2021. Shifting eye balance using monocularly directed attention in normal vision. J. Vis. 21, 4.

Wong, S.P., Hess, R.F., Mullen, K.T., 2025. Monocular eye-cueing shifts eye balance in amblyopia. J. Vis. 25, 6.

Yan, Y., Zhaoping, L., Li, W., 2018. Bottom-up saliency and top-down learning in the primary visual cortex of monkeys. Proc. Natl. Acad. Sci. U. S. A. 115, 10499–10504.

Zhang, P., Jiang, Y., He, S., 2012a. Voluntary attention modulates processing of eye-specific visual information. Psychol. Sci. 23, 254–260.

Zhang, X., Zhaoping, L., Zhou, T., Fang, F., 2012b. Neural activities in v1 create a bottom-up saliency map. Neuron 73, 183–192.

Zhaoping, L., 2008. Attention capture by eye of origin singletons even without awareness--a hallmark of a bottom-up saliency map in the primary visual cortex. J. Vis. 8, 1 1–18.

Zhaoping, L., 2012. Gaze capture by eye-of-origin singletons: interdependence with awareness. J. Vis. 12, 17.

Zhaoping, L., 2016. From the optic tectum to the primary visual cortex: migration through evolution of the saliency map for exogenous attentional guidance. Curr. Opin. Neurobiol. 40, 94–102.

Zhaoping, L., 2018. Ocularity Feature Contrast Attracts Attention Exogenously. Vision (Basel) 2.

Zhaoping, L., 2023. Peripheral and central sensation: multisensory orienting and recognition across species. Trends Cogn. Sci. 27, 539–552.

Zhaoping, L., Guyader, N., 2007. Interference with bottom-up feature detection by higher-level object recognition. Curr. Biol. 17, 26–31.

Zhaoping, L., Zhe, L., 2015. Primary Visual Cortex as a Saliency Map: A Parameter-Free Prediction and Its Test by Behavioral Data. PLoS Comput. Biol. 11, e1004375.

